# Calcineurin-responsive zinc finger 1 (Crz1) contributes to stress tolerance and virulence in the pathogenic fungus *Trichosporon asahii*

**DOI:** 10.1101/2025.07.02.662784

**Authors:** Yuta Shimizu, Yasuhiko Matsumoto, Takashi Sugita

## Abstract

The pathogenic fungus *Trichosporon asahii* causes severe invasive fungal infections in immunocompromised patients with neutropenia. In *Cryptococcus neoformans*, calcineurin-responsive zinc finger 1 (Crz1) functions as a transcription factor downstream of the calcineurin signaling pathway and regulates the expression of genes involved in stress resistance and virulence. In *T. asahii*, Cna1 and Cnb1, which are key components of the calcineurin pathway, contribute to various stress responses and virulence. The role of Crz1 in stress tolerance and virulence in *T. asahii*, however, has remained unclear. Here we demonstrate that a *crz1* gene-deficient *T. asahii* mutant exhibited increased sensitivity to cell wall and endoplasmic reticulum stress. The *crz1* gene-deficient mutant was sensitive to Congo red and tunicamycin, but not to dithiothreitol or sodium dodecyl sulfate. Moreover, the virulence of the *crz1* gene-deficient mutant in the silkworm infection model was reduced. These phenotypes of the *crz1* gene-deficient mutant were restored by reintroducing the *crz1* gene, confirming the association between Crz1 and these phenotypes. The half maximal lethal dose of the *cnb1* gene-deficient *T. asahii* mutant was higher than that of the *crz1* gene-deficient mutant. These results suggest that Crz1 plays a role in mediating the stress responses and virulence of *T. asahii*. The involvement of Crz1 in the virulence of *T. asahii* is small, however, compared to that of the calcineurin.

## Introduction

*Trichosporon asahii* is a dimorphic basidiomycete that is widely distributed in nature and is commonly isolated from soil, a variety of animal hosts, and the human gastrointestinal tract (1–6). *T. asahii* is an opportunistic pathogen that can cause severe invasive fungal infections, particularly in immunocompromised hosts such as patients with neutropenia (7–9). The mortality rate of invasive fungal infections caused by *T. asahii* is 70%–80% (10). Because echinocandin-class antifungal agents commonly used in the clinical treatment of fungal infections exhibit minimal antifungal activity against *T. asahii*, breakthrough infections may occur in patients undergoing treatment with micafungin (11, 12). Furthermore, clinical isolates of *T. asahii* resistant to antifungal agents such as amphotericin B and azoles have been isolated (13, 14). Therefore, revealing the molecular mechanisms of *T. asahii* infection may contribute to the development of effective prevention and treatment strategies (9).

Calcineurin-responsive zinc finger 1 (Crz1) is a transcription factor that contains a zinc finger DNA-binding domain (15). In *Cryptococcus neoformans*, a pathogenic basidiomycetous fungus closely related to *T. asahii*, Crz1 contributes to fungal virulence mediated by activation of the calcineurin signaling pathway (16, 17) . Crz1 is dephosphorylated by the calcineurin– calmodulin complex, allowing its translocation from the cytoplasm to the nucleus (17). Once in the nucleus, Crz1 regulates the expression of genes required for growth at 37°C and resistance to membrane and cell wall stress in *C. neoformans* (16, 17). Calcineurin is involved in the virulence of *T. asahii*, as well as in its resistance to high-temperature stress, membrane stress, cell wall stress, and endoplasmic reticulum (ER) stress (18). Whether Crz1 in *T. asahii* plays a role in stress resistance and virulence, however, has remained unknown.

In this study, we generated a *crz1* gene-deficient *T. asahii* mutant and examined the effects of the *crz1* gene on the stress response and virulence of *T. asahii*. The *crz1* gene-deficient mutant exhibited normal growth but showed increased sensitivity to tunicamycin and Congo red. Moreover, the virulence of the *crz1* gene-deficient mutant against silkworms was reduced compared to that of the parental strain. These findings suggest that Crz1 in *T. asahii* contributes to resistance against various stresses and plays a role in its virulence.

## Materials & Methods

### Reagents

Nourseothricin was purchased from JENA BIOSCIENCE GMBH (Dortmund, Germany). Hygromycin B was obtained from Tokyo Chemical Industry Co., Ltd. (Tokyo, Japan). Cefotaxime sodium, D-glucose, agar, NaCl, sorbitol, and dithiothreitol (DTT) were purchased from FUJIFILM Wako Pure Chemical Corporation (Osaka, Japan). Congo red was purchased from Sigma-Aldrich Co., LLC (St. Louis, MO, USA). G418 was obtained from Enzo Life Science, Inc. (Farmingdale, NY, USA). Hipolypeptone was purchased from NIHON PHARMACEUTICAL Co., LTD (Tokyo, Japan). Tunicamycin (TM) was purchased from Cayman Chemical Company (Ann Arbor, MI, USA). Sodium dodecyl sulfate (SDS) was obtained from NIPPON GENE Co., LTD. (Tokyo, Japan).

### T. asahii strains

*T. asahii* MPU129 Δ*ku70* was used as the parental strain (19) . The *crz1* gene-deficient mutant of *T. asahii* and the *crz1* gene-complemented strain in which the *crz1* gene was reintroduced into the *crz1* gene-deficient mutant were used. Strain information is provided in Table 1.

**Table 1.** Strains used in this study.

| *T. asahii* strains | Relevant genotype | Background | Reference |
| --- | --- | --- | --- |
| MPU129 $\Delta ku70$ (Parent strain) | *ku70::nptII* | MPU129 | Matsumoto Y., *et al.* (19) |
| $\Delta crz1$ | *ku70::nptII, crz1::NAT1* | MPU129 $\Delta ku70$ | This study |
| Crz1 (Revertant) | *ku70::nptII, NAT1::crz1, hph* | MPU129 $\Delta ku70 \Delta crz1$ | This study |

### *T. asahii* culture condition

*T. asahii* strains were cultured as described previously (19) . *T. asahii* strains were grown on Sabouraud dextrose agar (SDA; 1% hipolypeptone, 4% dextrose, and 1.5% agar) containing G418 (100 µg/mL) and cefotaxime (100 µg/mL) at 27°C for 1–2 days.

### Estimation of Crz1 in *T. asahii*

The Crz1 amino acid sequence of *Cryptococcus neoformans* was obtained from the National Center for Biotechnology Information (NCBI) (https://www.ncbi.nlm.nih.gov/). BLASTp was used to search for a homolog of *C. neoformans* Crz1 in *T. asahii* (https://blast.ncbi.nlm.nih.gov/Blast.cgi). Hypothetical protein A1Q1_05631, having the highest homology to *C. neoformans* Crz1, was identified as Crz1 of *T. asahii*.

### Homology analysis of Crz1 between *T. asahii* and other fungi

The Crz1 amino acid sequences of *T. asahii*, *Candida albicans*, *Aspergillus fumigatus*, *Cryptococcus neoformans*, and *Saccharomyces cerevisiae* were obtained from the NCBI. Homology analysis with these Crz1 proteins was performed using BLASTp. A phylogenetic tree was constructed following the maximum likelihood method using MEGA11 Software (https://www.megasoftware.net/).

### Construction of the *crz1* gene-deficient *T. asahii* mutant and the complement strain

The *crz1* gene-deficient *T. asahii* mutant and complement strain were constructed according to previous reports (18–21). Approximately 1000–1500 bp of the 5’-UTR and 3’-UTR of the *crz1* gene were cloned upstream and downstream of the *NAT1* gene region of the pUC19-NAT1 vector by the In-Fusion method (In-Fusion HD Cloning Kit, Takara Bio Inc., Shiga, Japan). The *crz1* gene-targeting DNA fragment 5’-UTR (*crz1*)-NAT1-3’UTR (*crz1*) was amplified by polymerase chain reaction (PCR) using the constructed plasmid. The *crz1* gene-targeting DNA fragment (2 µL) was introduced to competent cells (40 µL) of the *T. asahii* parent strain by electroporation. The electroporation condition used a time constant protocol (1800 V, 5 ms) with a 2-mm gap cuvette (Bio-Rad Laboratories, Inc., CA, USA) and Gene Pulser Xcell (Bio-Rad Laboratories, Inc., CA, USA). The electroporated cells were then incubated in 500 µL of YPD medium containing 0.6 M sorbitol at 27°C for 3 h. After incubation, the cells were spread onto SDA containing nourseothricin (300 µg/mL) and cultured at 27°C for 3–5 days. Candidate colonies grown on the drug-containing SDA were isolated, and their genomes were extracted to confirm *crz1* gene deficiency by PCR.

To construct the *crz1* gene-complemented strain, the *crz1* gene and *hph* gene conferring hygromycin B resistance were cloned between the downstream region of the *crz1* 5’-UTR and the upstream region of the 3’-UTR in the targeting plasmid used for constructing the crz1 gene deletion mutant. The DNA fragment 5’-UTR (*crz1*)-*crz1*-*hph*-3’UTR (*crz1*) was amplified by PCR using the constructed plasmid. Competent cells (40 µL) of the *crz1* gene-deficient mutant were mixed with 2 µL of the PCR-amplified DNA fragment (5’-UTR (*crz1*)-*crz1*-*hph*-3’UTR (*crz1*)). To introduce the DNA into the cells, electroporation was performed using the time constant protocol (1800 V, 5 ms). The cells were then suspended in 500 µL of YPD medium containing 0.6 M sorbitol and cultured at 27°C for 6 h. The cell suspension was spread onto YPD medium containing hygromycin B (200 µg/mL) and cultured at 27°C for 5 days. Colonies that grew were isolated, and their genomes were extracted to confirm reintegration of the *crz1* gene and *hph* gene by PCR. Primers used in this study are shown in Table 2.

**Table 2.**
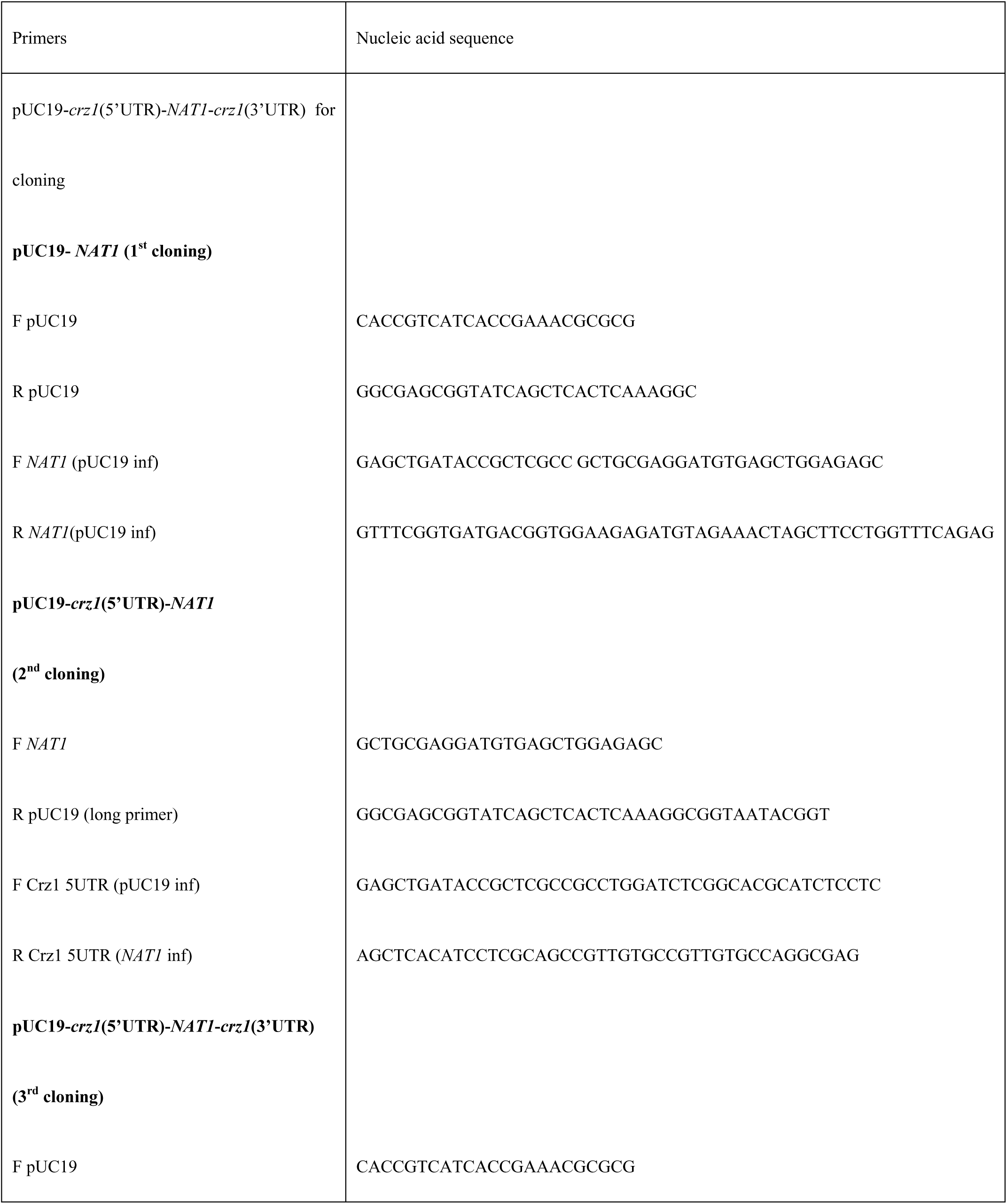
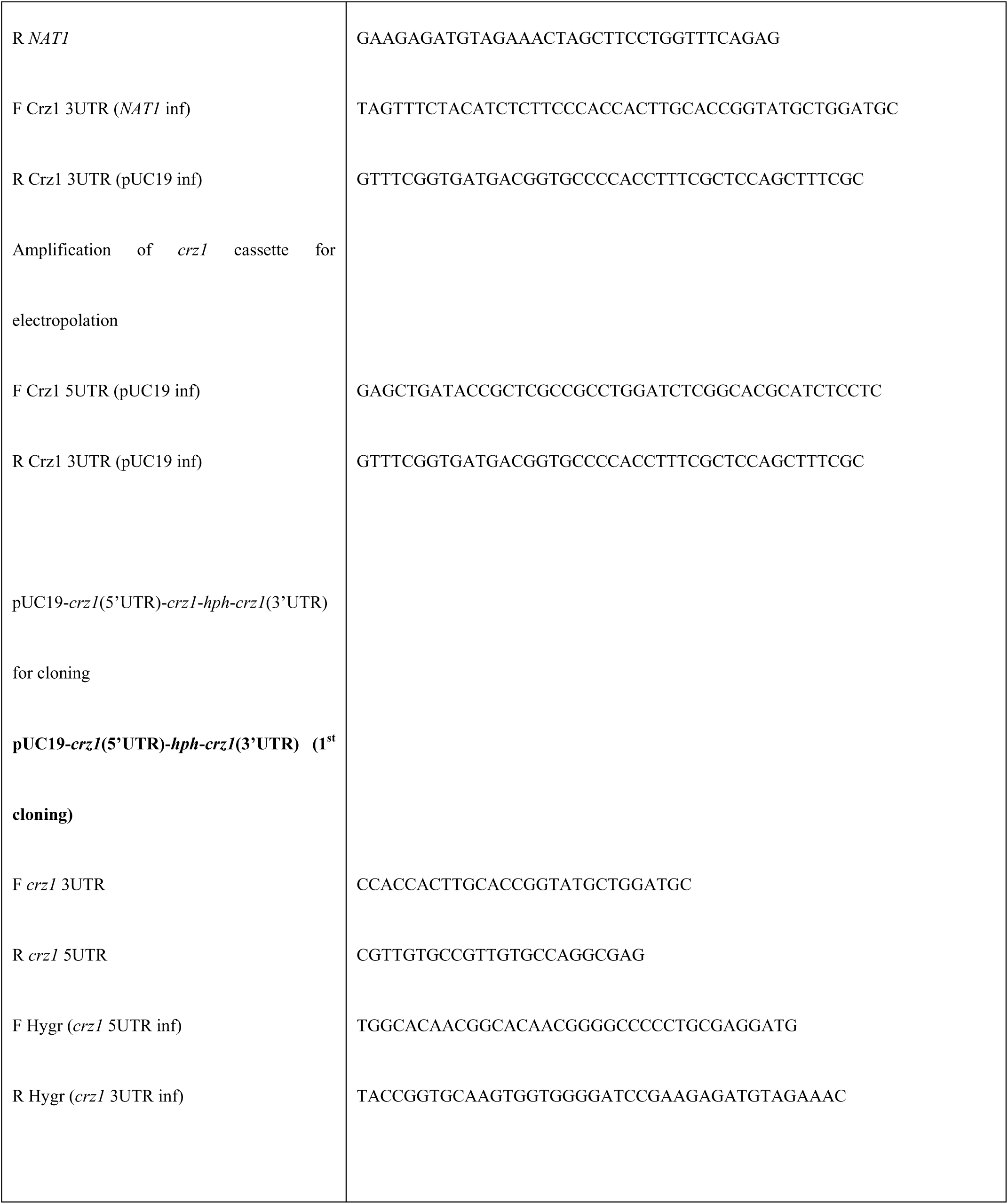
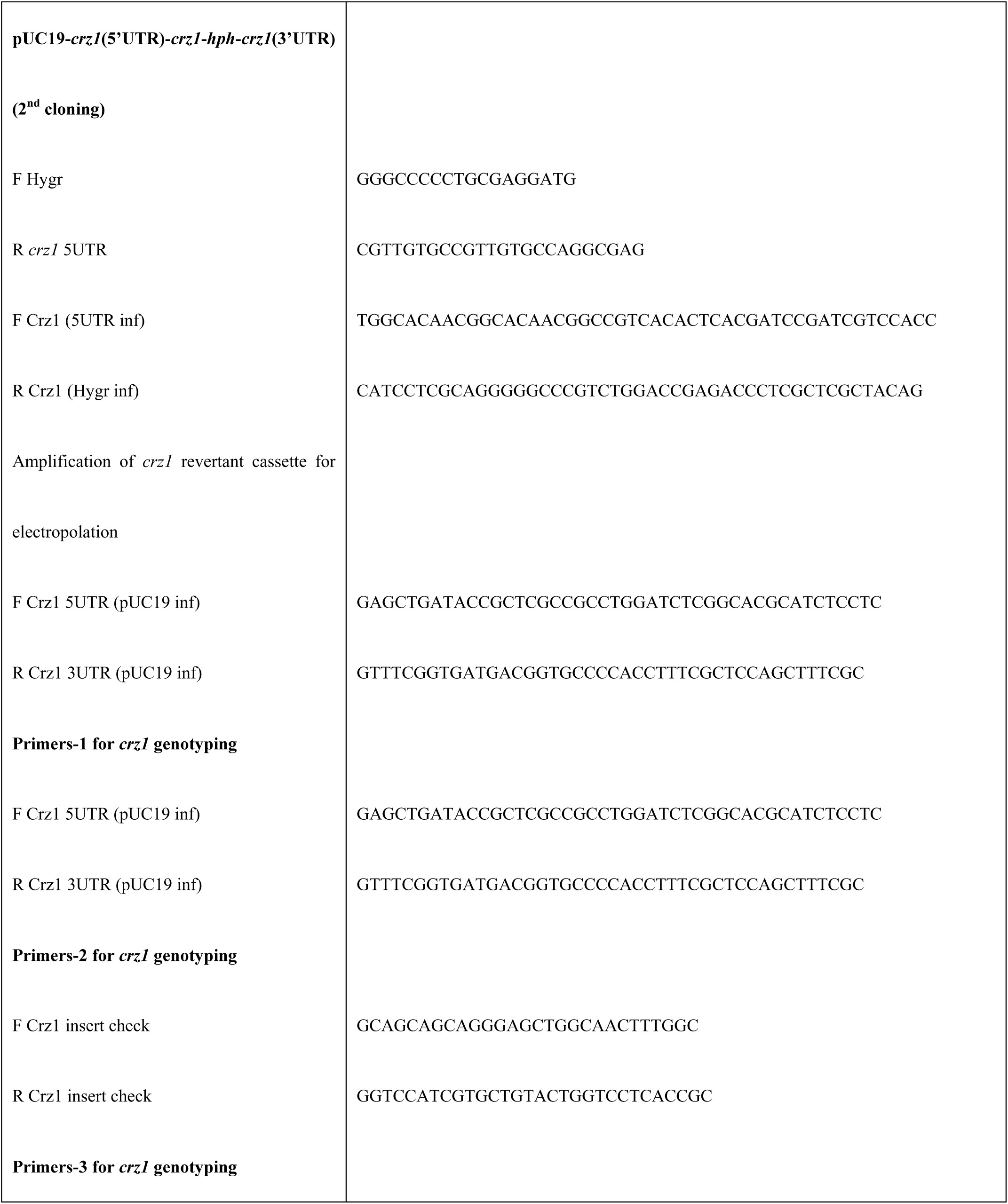
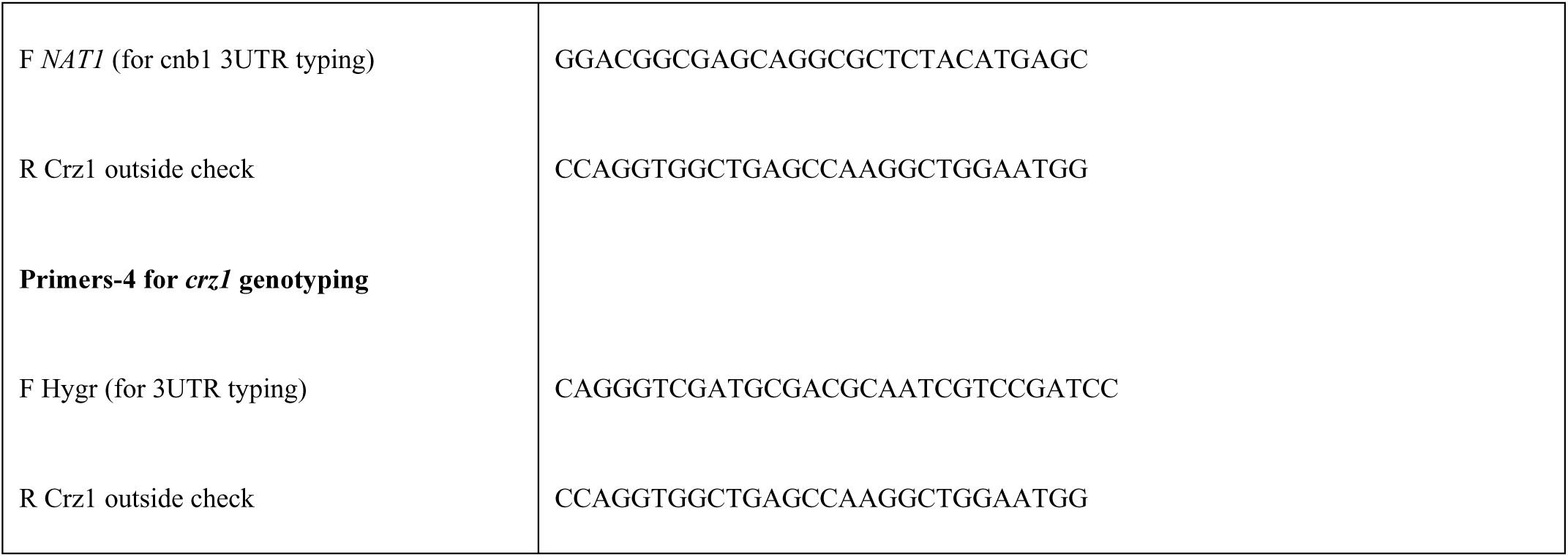
Primers used in this study.

### Stress sensitivity test

Stress sensitivity tests were performed according to the previous report (18, 20). *T. asahii* strains were cultured on SDA containing cefotaxime (100 µg/mL) at 27°C for 1 day. *T. asahii* cells were suspended in saline, and the cell suspension was adjusted to an absorbance of 1 at 630 nm. Ten-fold serial dilutions of each cell suspension were prepared using saline. Cell suspension (5 µL) was spotted onto SDA containing SDS, Congo red, TM, and DTT. Photographs were taken after cultivation at 27°C for 24–72 h.

### Silkworm infection experiment

Silkworm infection experiments were performed as described previously (18, 20, 21). *T. asahii* strains were grown on YPD agar containing cefotaxime (100 µg/mL) at 27°C for 1 day. *T. asahii* cells were suspended in saline and filtered through a cell strainer with a 40-µm pore size. The absorbance of the pass-through *T. asahii* cells at 630 nm was adjusted to 0.4. The *T. asahii* cell suspension was diluted 400-fold with saline, and 50 µL of the diluted suspension was injected into the silkworm hemolymph from the dorsal side using a 1-mL tuberculin syringe (Terumo Medical Corporation, Tokyo, Japan). The infected silkworms were fed an artificial diet (Silkmate 2S) and maintained at 37°C with monitoring for 4 days.

The half maximal lethal dose (LD_50_) of *T. asahii*, the dose required to kill half of the silkworms, was determined by modifying a previously reported method (20) . *T. asahii* (1.6 × 10^1^ to 1.5 × 10^7^ cells/50 µL) was injected into the silkworm hemolymph, and the silkworms were reared at 37°C with Silkmate 2S. The number of surviving silkworms (n = 4 per group) was measured 72 h post-injection. The LD_50_ was determined using a simple logistic regression model based on curves obtained from three independent experiments analyzed with Prism software (GraphPad Software, LLC, San Diego, CA, USA, https://www.graphpad.com/scientific-software/prism/).

### Statistical analysis

All experiments were performed at least three times, and representative results are shown. The statistical significance of differences in the growth of *T. asahii* between groups was calculated by Student’s *t*-test. The significance of differences between groups in the silkworm infection experiment was calculated by the log-rank test based on graphs constructed using the Kaplan-Meier method with JMP Pro 17 (https://www.jmp.com/ja_jp/support/jmp-software-updates.html). P < 0.05 was considered statistically significant.

## Results

### Identification of Crz1 protein in *T. asahii* and its conservation among fungi

We estimated *T. asahii* Crz1 based on *in silico* homology analysis. In *C. neoformans*, Crz1 is required for tolerance to various stressors as well as for virulence (22). A hypothetical protein of *T. asahii* (A1Q1_05631) was estimated as *T. asahii* Crz1. The percent identity and query cover of *T. asahii* A1Q1_05631 to the Crz1 protein of *C. neoformans* were 48.8% and 86%, respectively (Table 3). Therefore, we estimated that hypothetical protein A1Q1_05631 was *T. asahii* Crz1. Homology analysis of the amino acid sequence of the *T. asahii* Crz1 protein was conducted based on the genomic information of representative model fungi (*Saccharomyces cerevisiae*) and pathogenic fungi (*Cryptococcus neoformans*, *Candida albicans*, and *Aspergillus fumigatus*). The *T. asahii* Crz1 protein exhibited amino acid sequence identities of 49%, 48.8%, 42.1%, and 46% with the Crz1 proteins of *S. cerevisiae*, *C. neoformans*, *C. albicans*, and *A. fumigatus*, respectively (Table 3). A phylogenetic tree constructed from the amino acid sequence data of these proteins revealed that, among these fungi, the *T. asahii* Crz1 protein was most similar to the *C. neoformans* Crz1 protein (Fig. 1A). The *T. asahii* Crz1 protein contained the functional zinc finger domain similar to other fungal Crz1 proteins (Fig. 1B). Furthermore, two cysteines and two histidines essential for activity within the zinc finger domain were conserved across all fungi examined fungi (Fig. 1C).

**Fig. 1.**
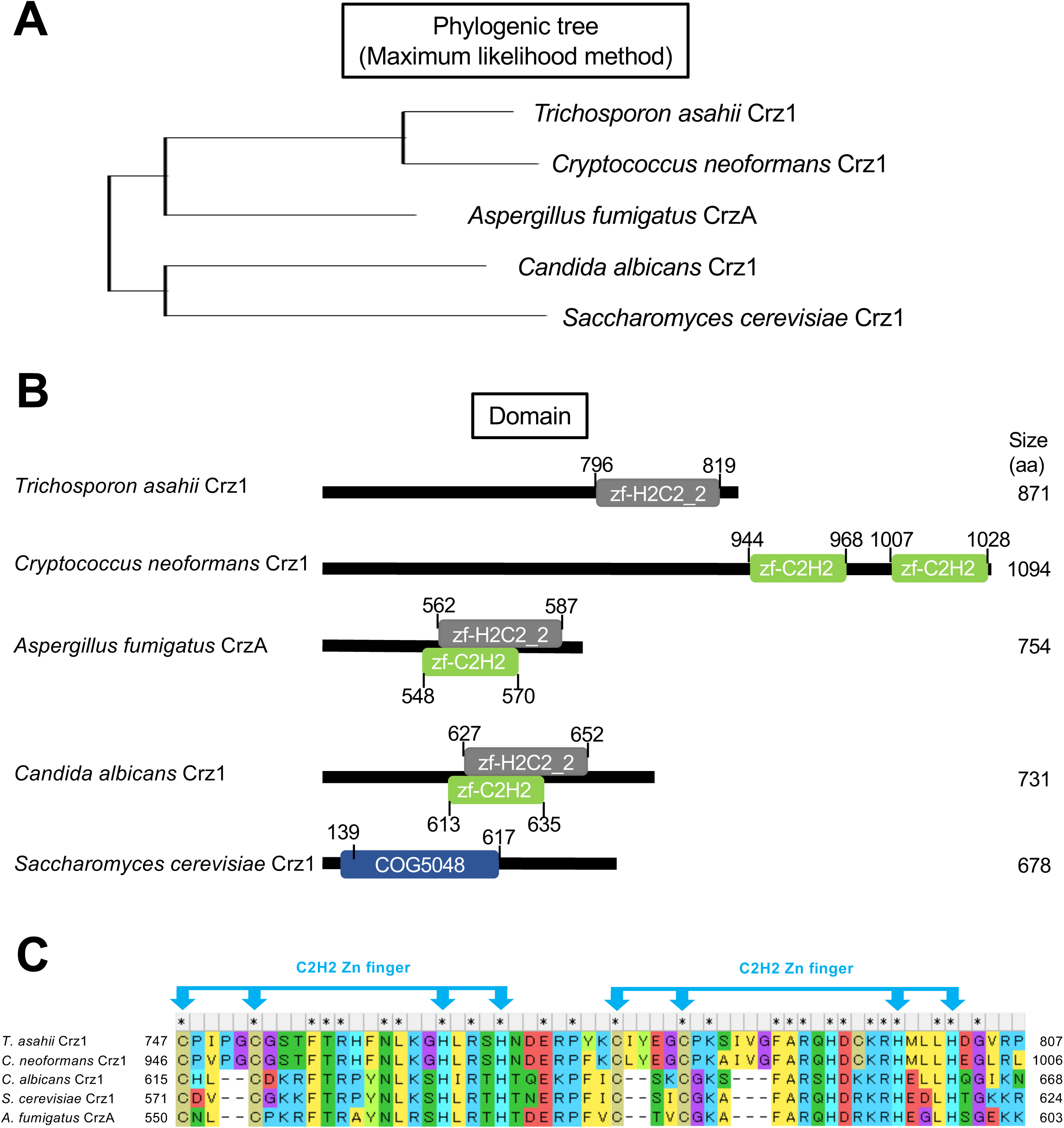
Homology analysis of Crz1 in *T. asahii* among various fungi. (A) Phylogenetic trees were constructed using the maximum likelihood method for Crz1 proteins of *T. asahii*, *C. neoformans*, *C. albicans*, *S. cerevisiae*, and CrzA of *A. fumigatus*. (B) Schematic representation of the domains presents in Crz1 and CrzA proteins. zf-H2C2_2, zf-C2H2, and COG5048 are zinc finger domains. (C) Amino acid sequences of the zinc finger domain in Crz1 and CrzA proteins. The zinc finger domain contains two C2H2 Zn fingers. Amino acid alignment analysis and domain prediction were performed using MEGA11 Software.

**Table 3.** Conservation of Crz1 among fungi including *T. asahii*.

| Species | Percent identify (%) | Query coverage (%) |
| --- | --- | --- |
| *T. asahii* | 100 | 100 |
| *C. neoformans* | 48.82 | 86 |
| *A. fumigatus* | 46.08 | 10 |
| *S. cerevisiae* | 49.07 | 11 |
| *C. albicans* | 42.11 | 13 |

### Construction of *crz1* gene-deficient *T. asahii* mutant

Next, we attempted to construct a *crz1* gene-deficient mutant of *T. asahii*. The DNA fragment required for *crz1* gene deficiency contained the *NAT1* gene, which confers resistance to nourseothricin (Fig. 2A). Insertion of this DNA fragment into the genome by homologous recombination confers resistance to nourseothricin (Fig. 2A). The DNA fragment was introduced into the *T. asahii* cells by electroporation, and transformants resistant to nourseothricin were obtained (Fig. 2B). Genomic DNA was extracted from the nourseothricin-resistant transformants, and *crz1* gene deficiency was confirmed by PCR (Fig. 2C, D). Moreover, we generated the *crz1* gene-complemented strain from the *crz1* gene-deficient mutant. The DNA fragment used for constructing the *crz1* gene-complemented strain contained the *hph* gene, which confers resistance to hygromycin B (Fig. 2A). This DNA fragment was introduced into cells of the *crz1* gene-deficient mutant via electroporation, resulting in transformants resistant to hygromycin B (Fig. 2B). Reintroduction of the *crz1* gene was confirmed by PCR using genomic DNA extracted from the hygromycin B-resistant transformants (Fig. 2C, D). These results suggest that the *crz1* gene-deficient mutant and the *crz1* gene-complemented strain of *T. asahii* were successfully generated.

**Fig. 2.**
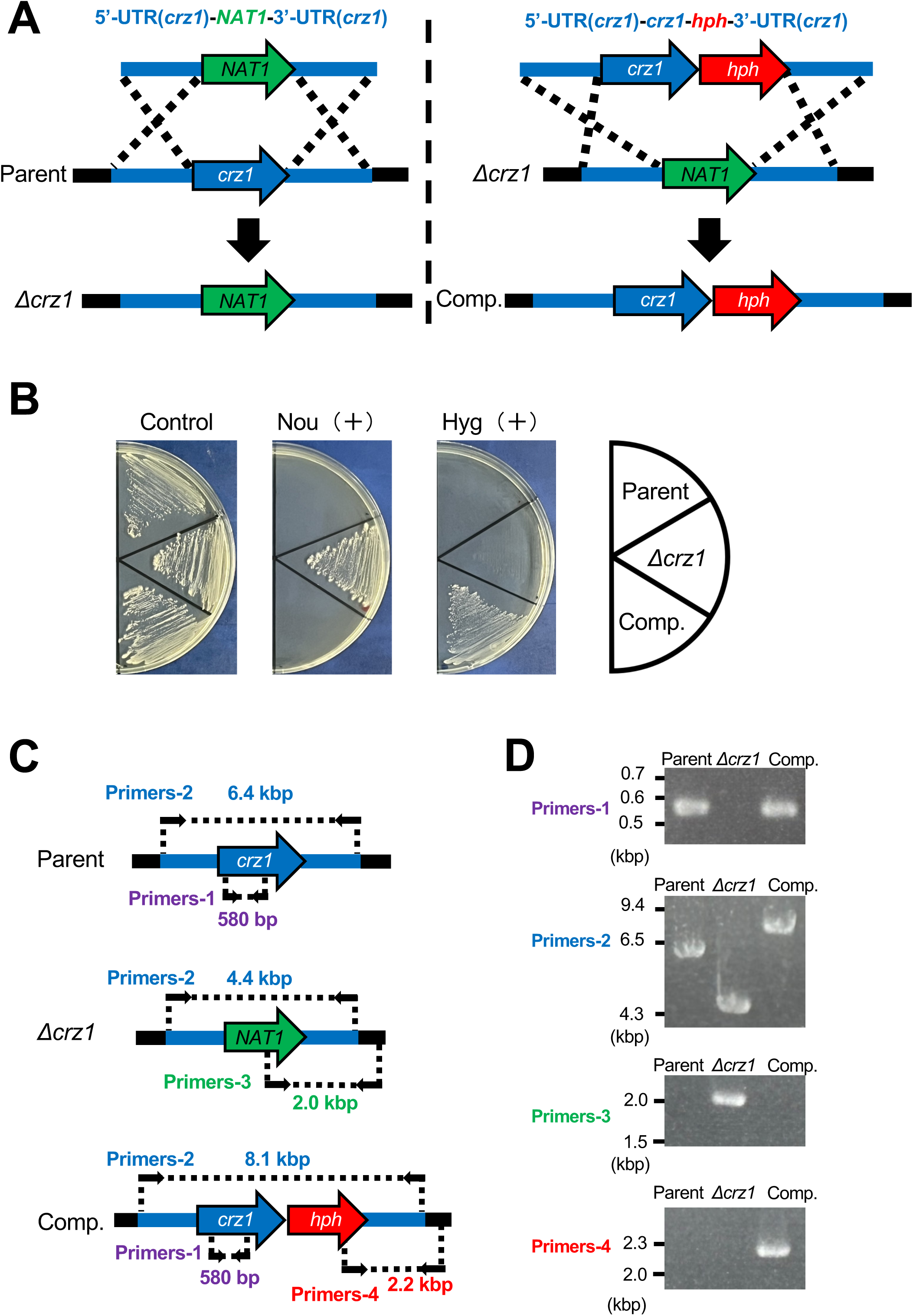
Establishment of *crz1* gene-deficient mutant and *crz1* gene-complemented *T. asahii* strains. (A) Schematic representation of the construction process for the *crz1* gene-deficient and the *crz1* gene-complemented mutant *T. asahii* strains. The genomic structures of the gene-deficient and complemented strains are also illustrated. (B) The parental strain (Parent), *crz1* gene-deficient strain (Δ*crz1*), and *crz1* gene-complemented strain (Comp.) were plated on SA medium containing nourseothricin (Nou; 100 µg/mL) or hygromycin B (Hyg B; 100 µg/mL) and incubated at 27°C for 2 days. (C) The positions of the primers and expected DNA fragment sizes are indicated to confirm the genomic structures of the *crz1* gene-deficient mutant and complemented strain by PCR using extracted genomic DNA. (D) DNA electrophoresis of PCR products was performed to confirm the genotypes of the *crz1* gene-deficient mutant (Δ*crz1*) and its complemented strain (Comp.).

### Role of the *crz1* gene under high-temperature stress

The *crz1* gene-deficient strain of *C. neoformans* exhibits delayed growth at 39°C (22). Calcineurin is involved in the growth of *T. asahii* at 40°C (18). We investigated whether the *crz1* gene contributes to high-temperature stress response in *T. asahii*. The *crz1* gene-deficient mutant of *T. asahii* did not exhibit delayed growth at 40°C compared to the parental strain (Fig. 3). This finding suggests that the *crz1* gene is not involved in the tolerance of *T. asahii* to high-temperature stress.

**Fig. 3.**
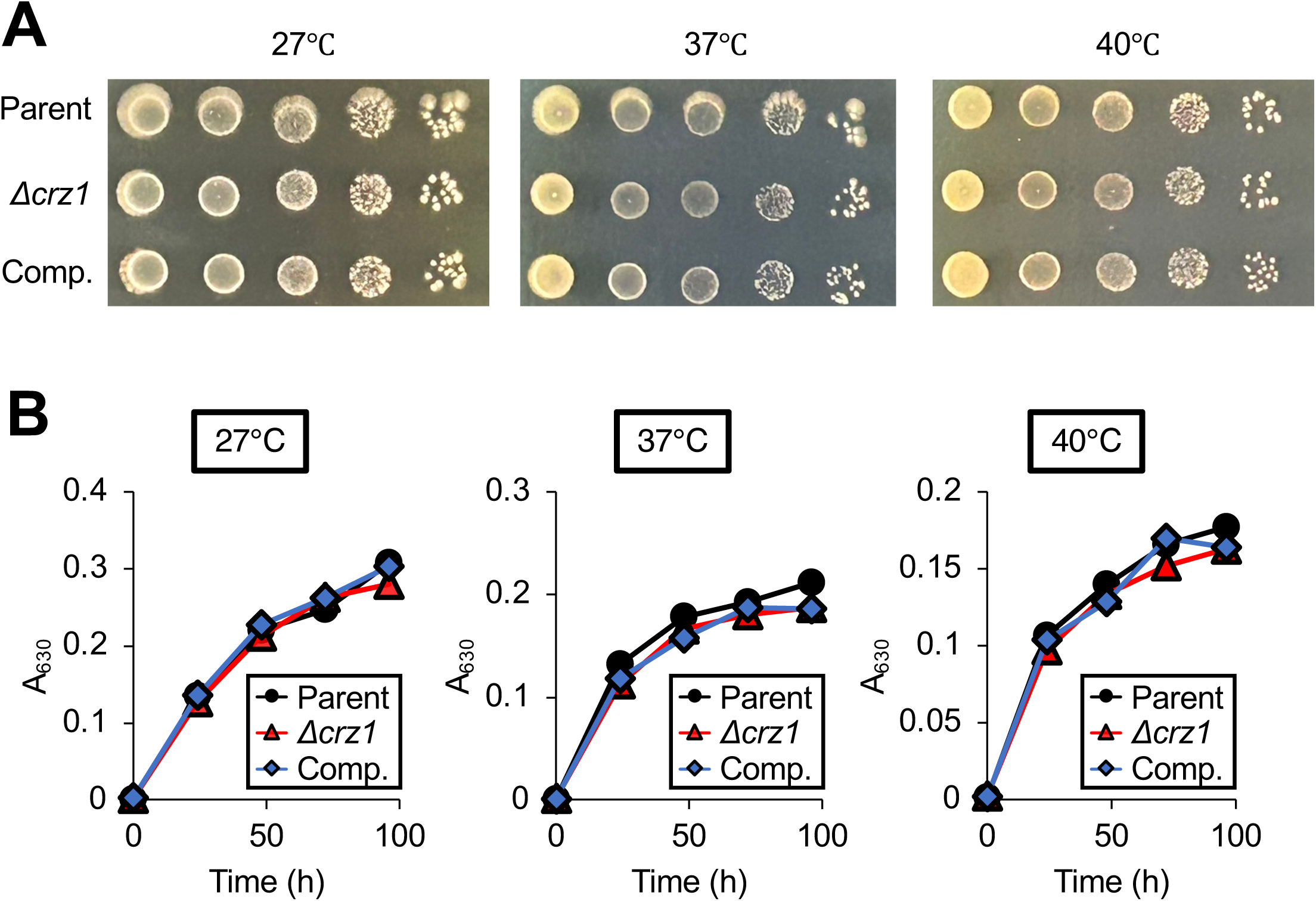
Temperature sensitivity of the *crz1* gene-deficient *T. asahii* mutant. (A) The parental (Parent), *crz1* gene-deficient mutant (Δ*crz1*), and *crz1* gene-complemented (Comp.) *T. asahii* strains were cultured on SA medium at 27°C for 2 days. *T. asahii* cells were suspended in saline to prepare a 10-fold serial dilution series. Five microliters of each cell suspension were spotted onto SA medium and incubated at 27°C, 37°C, or 40°C for 48 h. (B) The parental (Parent), *crz1* gene-deficient mutant (Δ*crz1*), and *crz1* gene-complemented (Comp.) *T. asahii* strains were cultured in Sabouraud dextrose liquid medium at 27°C, 37°C, or 40°C, and the optical density at 630 nm was measured.

### Role of the *crz1* gene in tolerance to various stresses

The *crz1* gene-deficient mutant of *C. neoformans* exhibits sensitivity to Congo red and SDS (18). Similarly, the *crz1* gene-deficient mutant of *C. albicans* shows sensitivity to SDS (23). Moreover, the *cnb1* gene-deficient mutant of *T. asahii* is sensitive to Congo red, SDS, TM, and DTT (18). We examined whether the *crz1* gene contributes to the response to various stressors. The *crz1* gene-deficient mutant of *T. asahii* exhibited delayed growth in the presence of Congo red, which induces cell wall stress, and TM, which induces ER stress, but not in the presence of SDS or DTT (Fig. 4). The *crz1* gene-complemented strain did not exhibit the phenotypes of cell wall stress and ER stress sensitivity observed in the *crz1* gene-deficient mutant (Fig. 4). Together, these findings suggest that the *T. asahii crz1* gene contributes to tolerance against cell wall stress and ER stress.

**Fig. 4.**
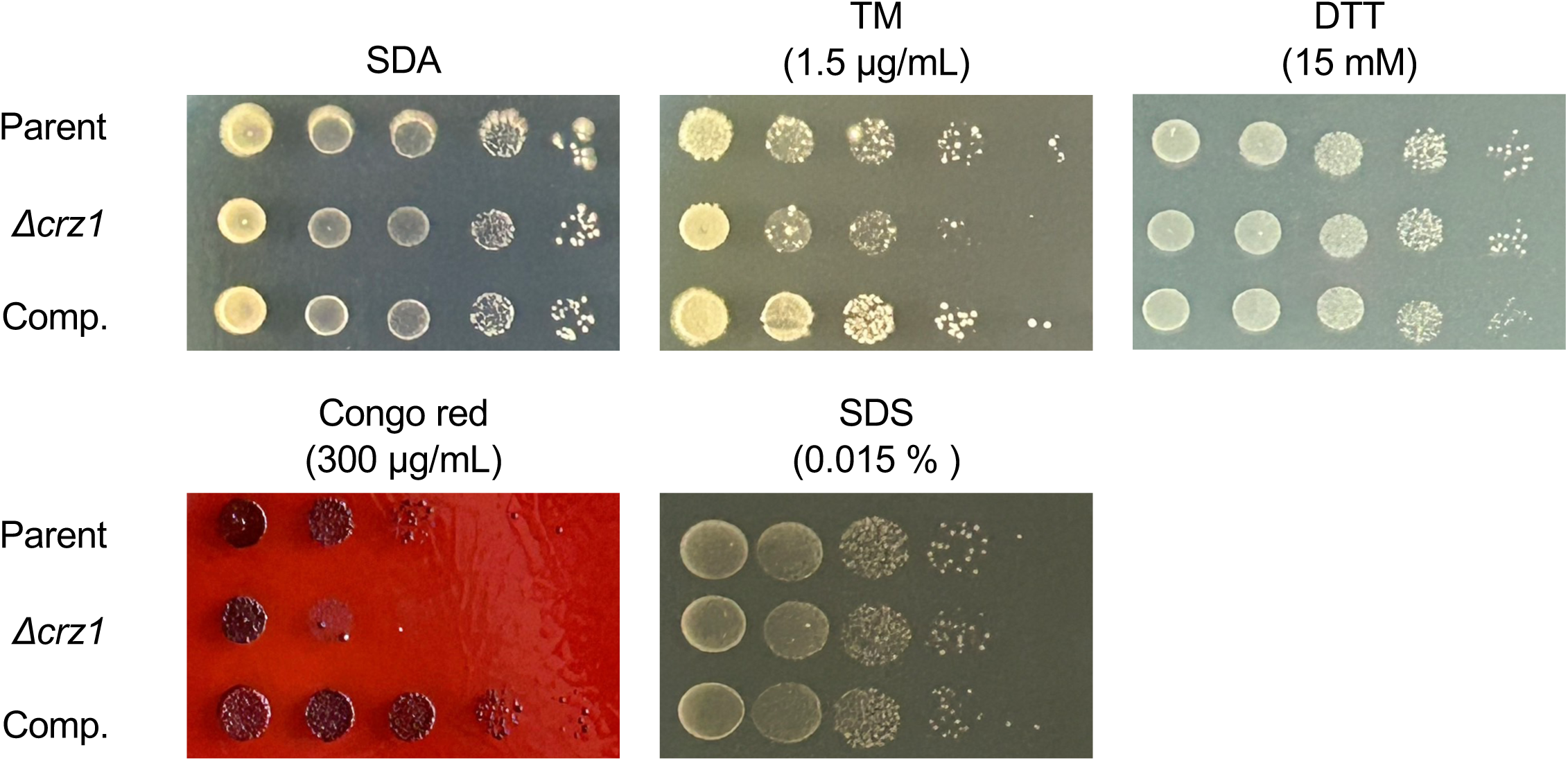
Stress sensitivity of the *crz1* gene-deficient *T. asahii* mutant. The parental (Parent), *crz1* gene-deficient mutant (Δ*crz1*), and *crz1* gene-complemented (Comp.) *T. asahii* strains were cultured on SA medium for 2 days. *T. asahii* cells were suspended in saline to prepare a 10-fold serial dilution series. The cell suspensions were spotted onto SDA medium containing either tunicamycin (TM) (1.5 µg/mL), dithiothreitol (DTT) (15 mM), Congo red (300 µg/mL), or sodium dodecyl sulfate (SDS) (0.015%) and incubated at 37°C for 72 h.

### Decreased virulence of the *crz1* gene-deficient mutant in a silkworm infection model

Crz1 in *C. neoformans*, *C. albicans*, and *A. fumigatus* is involved in its pathogenicity in mice (22, 24, 25). Therefore, we used a silkworm infection model to investigate whether *crz1* gene deficiency affects the virulence of *T. asahii*. Silkworms injected with the *crz1* gene-deficient mutant survived longer than those injected with the parental strain (Fig. 5A). The LD_50_ value of the *crz1* gene-deficient mutant was approximately five times higher than that of the parental strain (Fig. 5B and C). This phenotype was not observed in the complemented strain (Fig. 5). These findings suggest that the *crz1* gene contributes to the virulence of *T. asahii* against silkworms.

**Fig. 5.**
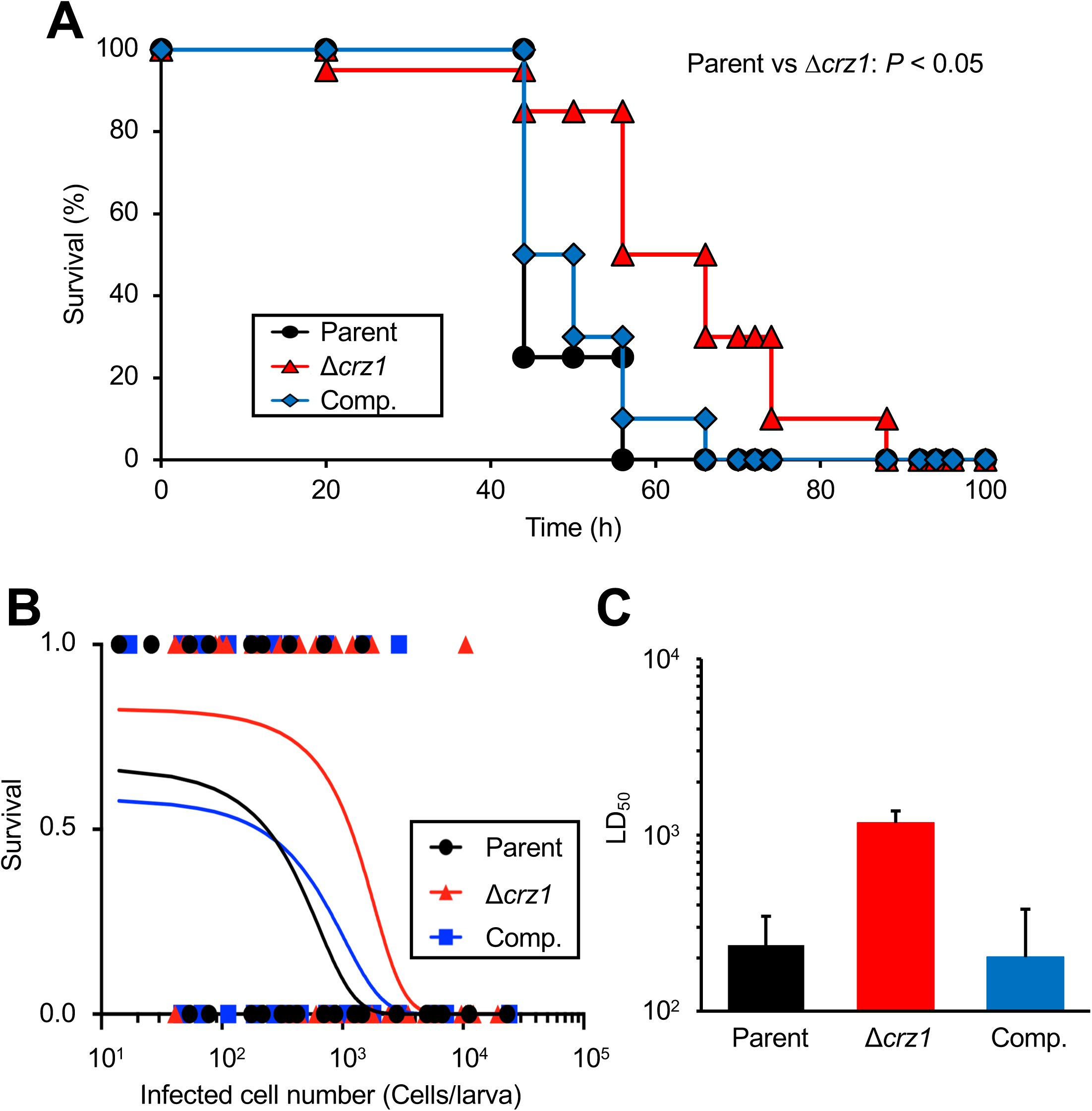
Virulence of the *crz1* gene-deficient *T. asahii* mutant against silkworms. (A) The parental (Parent; 160 cells/larva), *crz1* gene-deficient mutant (Δ*crz1*; 155 cells/larva), and *crz1* gene-complemented (Comp.; 210 cells/larva) *T. asahii* strains were injected into the hemocoel of silkworms, and their survival was monitored at 37°C for 96 h. The statistical significance of differences between the parental strain group and the *crz1* gene-deficient mutant group was determined by the log-rank test based on Kaplan-Meier survival curves. P-values < 0.05 were considered statistically significant. A total of 20 silkworms were used per group. (B) The survival rates of silkworms at 37°C were determined 72 h after *T. asahii* injection. Surviving and dead silkworms were scored as 1 and 0, respectively. A total of 48–64 silkworms were used per group. Curves were drawn by combining the results of three independent experiments using a simple logistic regression model. (C) The LD_50_ values were determined from the curves shown in panel B.

### Comparison of phenotypes between *crz1* and *cnb1* gene-deficient *T. asahii* mutants

In *C. neoformans*, Crz1 functions as a downstream factor of calcineurin (22). Next, we compared the phenotypes between *crz1* and *cnb1* gene-deficient *T. asahii* mutants. The *cnb1* gene-deficient mutant exhibited delayed growth at 40°C, whereas the *crz1* gene-deficient mutant did not show sensitivity at 40°C (Fig. 6A, B). The *cnb1* gene-deficient mutant was sensitive to TM, dithiothreitol DTT, Congo red, and SDS. The sensitivity of the *crz1* gene-deficient mutant to TM and Congo red was lower than of the *cnb1* gene-deficient mutant (Fig. 6C). While the *crz1* gene-deficient mutant exhibited reduced virulence in silkworms, the LD_50_ value of the *crz1* gene-deficient mutant in the silkworm infection system was 100-fold lower than that of the *cnb1* gene-deficient mutant (Fig. 6D-F). These results suggest that the phenotypes of the *crz1* gene-deficient *T. asahii* mutant were weaker than those of the *cnb1* gene-deficient *T. asahii* mutant.

**Fig. 6.**
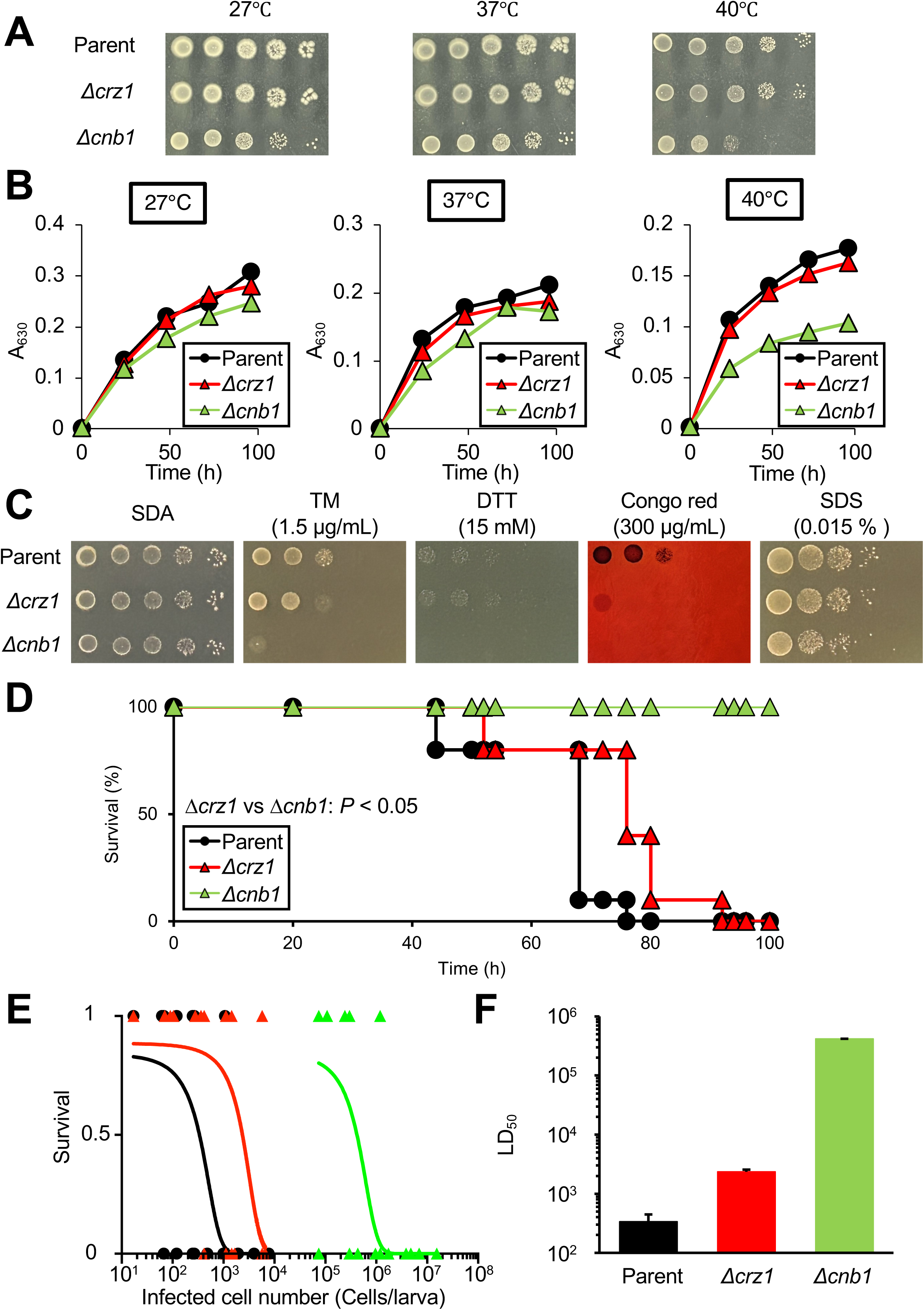
Comparison of the *crz1-* and *cnb1*-gene-deficient *T. asahii* mutant phenotypes. (A) The parental (Parent), *crz1* gene-deficient mutant (Δ*crz1*), and *cnb1* gene-deficient mutant (Δ*cnb1*) *T. asahii* strains were cultured on SDA at 27°C for 2 days. *T. asahii* cells were suspended in saline to prepare a 10-fold serial dilution series. Five microliters of each cell suspension was spotted onto SA medium and incubated at 27°C, 37°C, or 40°C for 48 h. (B) The parental (Parent), *crz1* gene-deficient mutant (Δ*crz1*), and *cnb1* gene-deficient mutant (Δ*cnb1*) *T. asahii* strains were cultured in Sabouraud dextrose liquid medium at 27°C, 37°C, or 40°C, and the optical density at 630 nm was measured. (C) The parental (Parent), *crz1* gene-deficient mutant (Δ*crz1*), and *cnb1* gene-deficient mutant (Δ*cnb1*) *T. asahii* strains were cultured on SDA for 2 days. *T. asahii* cells were suspended in saline to prepare a 10-fold serial dilution series. The cell suspensions were spotted onto SDA containing either tunicamycin (TM) (1.5 µg/mL), dithiothreitol (DTT) (15 mM), Congo red (300 µg/mL), or sodium dodecyl sulfate (SDS) (0.015%) and incubated at 37°C for 72 h. (D) The parental (Parent; 278 cells/larva), *crz1* gene-deficient mutant (Δ*crz1*; 298 cells/larva), and *cnb1* gene-deficient mutant (Δ*cnb1*; 400 cells/larva) *T. asahii* strains were injected into the silkworm hemolymph, and their survival was monitored at 37°C for 96 h. The statistical significance of differences between the parental strain group and the *crz1* gene-deficient mutant group was determined by the log-rank test based on Kaplan-Meier survival curves. P-values < 0.05 were considered statistically significant. Each group contained 20 silkworms. (E) Survival rates of silkworms at 37°C were determined 72 h after *T. asahii* injection. Surviving and dead silkworms were scored as 1 and 0, respectively. Each group contained 48–64 silkworms. Curves were drawn by combining the results of three independent experiments using a simple logistic regression model. (F) The LD_50_ values were determined from the curves shown in panel E.

## Discussion

In this study, we generated a *T. asahii crz1* gene-deficient mutant and demonstrated that the mutant exhibited reduced tolerance to several stressors as well as reduced virulence in the silkworm infection model. These findings suggest that both virulence and stress resistance are regulated through Crz1-dependent pathways in *T. asahii*.

The *T. asahii crz1* gene contributes to its stress responses against TM and Congo red, but not against high temperature, DTT, or SDS. TM induces ER stress (26) and Congo red induces cell wall stress (27). Therefore, Crz1 might regulate ER and cell wall stress tolerance in *T. asahii*. In *C. neoformans*, Crz1 has crucial roles in its tolerance to high temperature, TM, Congo red, DTT, and SDS (22, 28–31). Moreover, *crz1* gene deficiency leads to increased sensitivity to TM and SDS in *C. albicans* (23, 32, 33). Therefore, Crz1 proteins in *T. asahii*, *C. neoformans*, and *C. albicans* commonly contribute to the response to ER stress induced by TM. On the other hand, the *crz1* gene is required for SDS tolerance in *C. neoformans* and *C. albicans*, but not in *T. asahii*. In *C. neoformans,* but not *T. asahii,* high temperature sensitivity is increased by *crz1* gene deficiency. The phenotypic differences observed among pathogenic fungi might be due to species-specific regulatory mechanisms involving *crz1*. Further studies are needed to compare how Crz1 regulates gene expression across different fungal species.

The *crz1* gene contributes to the virulence of *T. asahii* against silkworms. In *C. neoformans*, *C. albicans*, and *A. fumigatus*, Crz1 is required for their virulence in mouse infection experiments (22, 24, 25). Therefore, Crz1 regulates the expression of genes required for virulence in various pathogenic fungi, including *T. asahii*. The calcineurin and Hog1 signaling pathways have important roles in *T. asahii* virulence (18, 20). Therefore, understanding the relationships between Crz1 and other signaling pathways in *T. asahii* virulence is an important goal.

The phenotypes of the *crz1* gene-deficient mutant were weaker than those of the *cnb1* gene-deficient mutant, which is a calcineurin-deficient mutant. Crz1 is regulated by calcineurin through dephosphorylation (34). Therefore, the phenotypes of the *crz1* gene-deficient mutant and the *cnb1* gene-deficient mutant were compared. The *cnb1* gene-deficient *T. asahii* mutant exhibited sensitivity to high temperature, TM, DTT, Congo red, and SDS (16, 17). Growth of the *cnb1* gene-deficient mutant was delayed under exposure to high temperature, TM, DTT, Congo red, or SDS, in comparison with the *crz1* gene-deficient mutant. Moreover, the LD_50_ value of the *cnb1* gene-deficient mutant was more than 100-fold higher than that of the *crz1* gene-deficient mutant. Therefore, the contribution of Crz1 to tolerance to various stressors and to the virulence of *T. asahii* was small compared with that of calcineurin. We hypothesize that calcineurin regulates *T. asahii* virulence via Crz1-dependent and independent mechanisms. Further investigation into the roles of calcineurin and Crz1 in *T. asahii* virulence will be an important focus of future research.

In conclusion, Crz1 mediates tolerance to various stressors as well as virulence in *T. asahii*, but its contribution is smaller than that of calcineurin. Therefore, transcription factors other than Crz1 may act downstream of calcineurin.

## Acknowledgements

We thank Renta Endo, Momoka Matsumura, and Tomoya Sanbongi (Meiji Pharmaceutical University) for technical assistance rearing the silkworms. We also thank the editors of SciTechEdit International LLC (Highlands Ranch, CO, USA) for providing English-language editing. This study was supported by JSPS KAKENHI grant number JP23K06141 (Scientific Research (C) to Y.M.), in part by the Research Program on Emerging and Re-emerging Infectious Diseases of the Japan Agency for Medical Research and Development, AMED (Grant number JP23fk0108679h0401 to T.S.), and the Research and implementation promotion program through open innovation grants(Grant# JPJ011937, to the consortium where Y.M. serves as a member) from the Project of the Bio-oriented Technology Research Advancement Institution (BRAIN).

